# Assessing diatom-associated bacteria by a specific CARD-FISH protocol

**DOI:** 10.1101/2023.05.12.540504

**Authors:** Tran Quoc Den, Meinhard Simon

## Abstract

The cell surfaces of phytoplankton algae including diatoms are usually colonized by specific bacterial populations which interact with and affect growth of the host species. Catalyzed Reported Deposition Fluorescence in situ Hybridizazion (CARD-FISH) is a well-suited technique to visualize and identify algal-associated bacterial cells. Autofluorescence and the strongly structured cell surface of the algal cell make it difficult to quantify distinct populations of the colonizing bacterial communities. To overcome these limitations, we adopted a CARD-FISH method to this specific application by reducing the algal autofluorescence by an extra ethanol treatment and by stacking epifluorescence micrographs taken at different focal planes and merging them into a composite image. Cells of the diatom *Thalassiosira rotula* were used as host and incubated with a consortium of different bacterial strains and a natural bacterial community. Samples were concentrated either by filtration onto polycarbonate membranes or by centrifugation and analyzed with probes CF319a, GAM42a and ROS536. The results showed easily quantifiable bacterial cells and did not reveal any significant differences in the abundance of diatom-associated bacterial populations assessed by both methods. Our CARD-FISH protocol persuasively demonstrated that bacterial populations can be easily and reliably enumerated on diatom cells and presumably also on other algal cells and autotrophic biofilms.

## Introduction

Phytoplankton and heterotrophic bacterioplankton communities play a major role in the cycling of carbon, nitrogen and other elements in oceanic, coastal and freshwater ecosystems (Nelson et al. 1995, Falkowski 2012, Baltar et al. 2015, Okazaki et al. 2017, Pajares and Ramos 2019, Borics et al. 2021). The relationship and mutual interactions between these two groups of microorganisms have important implications and consequences for the function of these ecosystems. Microalgae such as diatoms and dinoflagellates provide heterotrophic bacteria with substrates such as carbohydrates and amino acids for growth (Durkin et al. 2009, Bennke et al. 2013, Amin et al. 2015, Cruz-López et al. 2018, Becker et al. 2020, Shibl et al. 2020, Vidal-Melgosa et al. 2021), whereas bacteria provide essential growth factors for the algal proliferation such as hormones, vitamins B_1_ and B_12_ (Croft et al. 2005, Grant et al. 2014, Amin et al. 2015, Cruz-López and Maske 2016, Cruz-López et al. 2018). Additionally, distinct bacteria, in particular members of *Bacteroidetes*, *Gammaproteobacteria* and the *Roseobacter* group, colonize and attach to the algal host cell, presumably enabled by encoded gene clusters for a tight attachment (Bauer et al. 2006, de Bentzmann et al. 2006, González et al. 2008, Bakenhus et al. 2018, Isaac et al. 2021). These physical interactions between heterotrophic bacteria and phytoplankton species have received much attention in the recent past because they can affect the development of either or both partners negatively or positively with ecosystem-wide consequences (Li et al. 2016, Cooper et al. 2019). The most suitable means to investigate these species-specific physical interactions is CARD-FISH and epifluorescence microscopy or Confocal Laser Scanning Microscopy (CLSM) (Bennke et al. 2013, Tran et al. 2022).

CARD-FISH (known also as tyramide signal amplification [TSA]-FISH) was introduced to overcome shortcomings of the original FISH method (DeLong et al. 1989), mainly low signal intensity (Schönhuber et al. 1997). Due to the remarkable effectiveness of this method, it has continuously been improved and adapted for investigating prokaryotic communities in different habitats, such as soil (Chatzinotas et al. 1998, Ferrari et al. 2006), sediment (Ishii et al. 2004, Ramm et al. 2012, Tischer et al. 2012), the marine pelagic zone (Pernthaler et al. 2002, Herndl et al. 2005, Bakenhus et al. 2017), marine macroalgal surfaces (Tujula et al. 2006, Brunet et al. 2021), freshwater (Sekar et al. 2003), wastewater (Pavlekovic et al. 2009) and protist grazing (Jezbera et al. 2005, Ballen-Segura et al. 2016, 2018, Piwosz et al. 2021). A few studies have investigated bacteria colonizing marine diatom aggregates using CARD-FISH (Bennke et al. 2013, Arandia-Gorostidi et al. 2022). Problems applying this method to bacteria colonizing diatoms, but also other microalgae, is the intense autofluorescence of the algae and to obtain high quality photos of the bacteria colonizing the three dimensional algal cells in different focal planes. The application of appropriate filter sets separating the excitation wavelength of the fluorescent dye of the probe and chlorophyll *a* (Arandia-Gorostidi et al. 2022) and the complete removal of cellular pigments can circumvent the problems with autofluorescence (Wada et al. 2016, Castillo et al. 2020). It is desirable for investigating bacterial populations attached to diatom and other microalgal cells to apply a method with a low background autofluorescence of the algal cells. Further, the application of an image analysis system and stacked images to produce high quality photos of the colonizing bacteria is highly preferred when CLSM is not available.

To achieve these aims we used a method to investigate bacterial populations colonizing diatom cells with a low autofluorescence and applied a program to stack images to generate high quality micrographs of the bacteria colonizing diatom cells. As the algal aggregates need to be concentrated we further tested whether there is any difference in concentrating the aggregates suspended in the sample by filtration or centrifugation. Consequently, the diatom *Thalassiosira rotula* was grown with a consortium of different bacterial strains affiliated to *Bacteroidetes*, *Gammaproteobacteria* and the *Roseobacter* group of *Alphaproteobacteria* and a natural bacterial community and subsampled for studying the attachment patterns of the bacteria.

## Methods

### Co-cultures of bacteria and the diatom *Thalassiosira rotula*

In order to determine and quantify attached bacteria on the host diatom, the axenic diatom *Thalassiosira rotula* was grown in enriched artificial seawater medium (ESAW, (Harrison et al. 1980, Tran et al. 2022) at 15°C in a 12:12 h light:dark cycle (irradiance 35–38 μmol m^−2^ s^−1^, Neon tubes Biolux and Fluora, Osram, Munich, Germany) for five days to reach the beginning of the exponential phase.

A consortium of five bacterial strains including *Gramella forsetii* (*Bacteroidetes*), *Glaciecola* sp. (*Gammaproteobacterium*), *Colwellia* sp. M166 (*Gammaproteobacterium*), *Roseovarius* sp. M141 (*Roseobacter*) and *Pseudophaeobacter* sp. (*Roseobacter*) was incubated at 20^°^C for four days in ESAW amended with 50% Marine Broth (MB, Dicfo 2216, Becton Dickinson, Franklin Lakes, NJ, USA) and subsequently for four days in ESAW supplemented with 5 mM of glucose. In addition, to test whether there is a difference in the effectiveness of the modified CARD-FISH procedures on the quantification of attached bacteria between natural communities and defined co-cultures of bacterial strains, fresh seawater was collected from the North Sea coast (Neuharlingersiel, 53° 41’ 59.237” N 7° 42’ 12.557” E) on December 23^rd^, 2020 and prefiltered through a 3 µm polycarbonate membrane (Whatman, Maidstone, UK) to remove large particles and phytoplankton and to collect the free-living bacteria. The bacterial cells in the filtrate and in the species consortium were washed twice in sterile ESAW medium by centrifugation (10,000 g; 2 min) before co-culturing with 5 days old axenic *T. rotula*. The mixed cultures of the natural community and the species consortium and the diatom were incubated at 15°C in a 12:12 h light:dark cycle for 21 days.

### Sample collection and fixation

Samples of the two treatments were collected at days 10 and 21 and fixed with 37% formaldehyde solution (1 ml sample fixed with 30 µl 37% formaldehyde; final concentration of 1.11% [v/v]) for 1 h at room temperature. These samples were filtered onto 5.0 µm pore-size polycarbonate filters (diameter, 25 mm; Isopore, Sigma-Aldrich, Merck, Darmstadt, Germany) using a filter holder (1225 Sampling Manifold Chambers; Millipore, Merck). Before being removed from the filter holder, each filter was washed by adding 30–40 ml of 1× Phosphate Buffered Saline solution (PBS: 137 mM NaCl, 2.7 mM KCl, 10 mM Na_2_HPO_4_, 1.8 mM KH_2_PO_4_) into the chamber to remove not firmly attached bacteria (free-living cells) and algal threads settled on the frustules during filtration. The threads settled on the frustules’ surface needed to be removed as these structures were found to carry attached bacteria as well (Supporting Information Fig. S1; Tran *et al*. 2022a). All filter samples were stored at -20°C until further analysis by CARD-FISH. At day 21, algal cells with the attached bacteria in the treatment with the species consortium were also collected by a pipette from the formaldehyde-fixed suspended sample and transferred into 2 ml safe-lock tubes (Eppendorf, Wesseling-Berzdorf, Germany). To avoid an excessive fixation, which may reduce accessibility of the probe, these tubes were stored at 4°C and the CARD-FISH analysis was performed within three days.

### Examination of diatom-attached bacteria using filtered samples

The assessment of attached bacteria by CARD-FISH was carried out using sections of the 5 µm polycarbonate filter according to the protocol described previously (Pernthaler et al. 2002, Bakenhus et al. 2017) with a modification to reduce chlorophyll autofluorescence. Filter sections were firstly embedded in low-melting point agarose (0.1% [w/v]; Biozym Plaque Agarose; Biozyme, Hessisch Oldendorf, Germany) to prevent cell loss. The autofluorescence of chlorophyll was reduced by immersing the samples in 96% ethanol for 10 min at room temperature (RT). For permeabilization, samples were treated with 10 mg ml^-1^ lysozyme (Thermo Fisher, Leiden, The Netherlands) solution for 45 min at 37°C. Inactivation of endogenous peroxidase was performed by 0.3% H_2_O_2_ solution for 10 min at RT. The hybridizations were carried out in glass humidity chambers at 46°C for 120 min using Horseradish Peroxidase (HRP)-labelled probes (Table 1). Probes in aliquots (50 ng µl^-1^) were diluted in hybridization buffer (35% formamide [FA] concentration) to obtain 0.5 and 1.0 ng µl^-1^ hybridization solution for the analysis. Hybridized cells on filters were washed with a washing buffer (1 M Tris–HCl, 0.5 M EDTA, 5 M NaCl, and 20% SDS) for 10 min at 48°C. The washing buffer was preadjusted to this temperature. The amplification stage was done using 1 µg ml^-1^ FITC-tyramide (fluorescein-5-isothiocyanate –tyramide; Sigma-Aldrich) solution for 30 min at 37°C. Samples were counterstained with a DAPI (4′,6-diamidino-2-phenylindole) solution (1 μg ml^-1^) for 3 min at RT. Finally, filter sections were embedded on slides using Citifluor and Vectashield (4:1 [v/v]). Samples after CARD-FISH and DAPI staining were stored in the dark at -20°C until examination by epifluorescence microscopy. All analyses were done in triplicate.

**Table 1.**
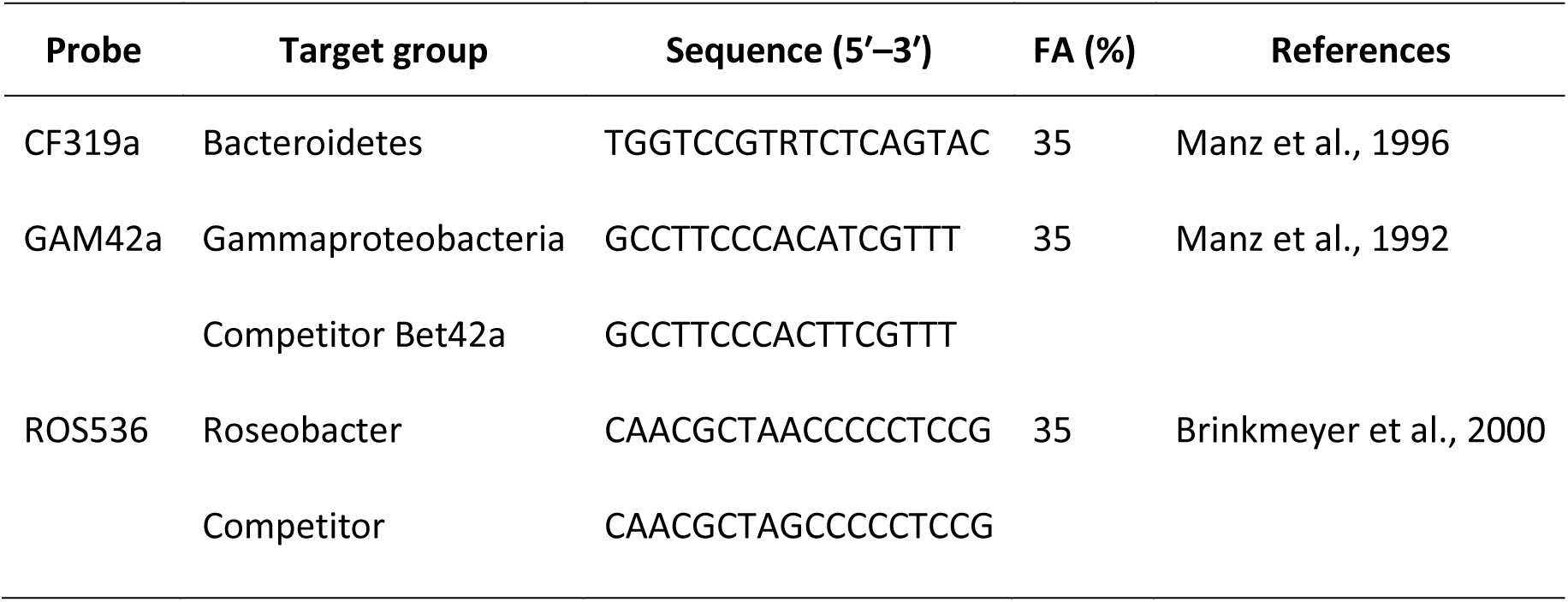
Target groups, sequences, % Formaldehyde (FA) and references of the HRP-probes used.

The diatom-attached bacterial cells were detected and quantified using a Zeiss Microscope AxioImager.Z2m (Carl Zeiss, Jena, Germany) including the software package AxioVision (version 40 V4.8.2.) with the objective 63x NA 1.46 (α Plan Apochromat objective) and the filter sets 01 for DAPI and 09 for FITC-tyramide, respectively. Fluorescence images were taken using the sensitive imaging black and white AxioCam MRm camera (Carl Zeiss). Because the visualization of algal thread structures is not possible with regular epifluorescence microscopy, in this study bacteria were considered as cells attached to the diatom when they were found to colonize directly the frustule or adhere to the side of the frustule of *T. rotula*. As a consequence, the frustule was used to normalize the bacterial cell abundance as cells frustule^-1^ on the host diatom.

On every algal frustule, DAPI-signals of total bacteria and FITC-signals of the attached bacteria were assessed. If numbers of attached bacteria were low, i.e. <10 cells frustule^-1^, and manually countable, the focal plane of the microscope was manually adjusted so that all attached bacterial cells were enumerated, also when located at different depths of the frustule. At higher bacterial abundances, signals of cells in both image channels in a field of view (FOV) were recorded in a series of micrographs taken at 2 - 5 focal planes. Afterwards, a set of images obtained at the same FOV of each channel was stacked using Picolay (www.picolay.de) and the enumeration of bacterial numbers was carried out manually or using the automated image analysis software ACMEtool3 (Bennke et al. 2016). The abundance of diatom-attached bacteria was acquired on the basis of cell numbers counted from 50 to 100 randomly selected diatom frustules.

After DAPI counterstaining the large nucleus of *T. rotula* cells covers a substantial area of the cell by its DNA fluorescence where DAPI signals of the attached bacteria are masked (Supporting Information Fig. S2). To circumvent this bias for assessing the numbers of attached bacteria, the absolute abundance of total attached bacteria detected with DAPI was calculated according to the equation N = 2 × (n + (*c* × n)) where N is the number of attached bacteria on one diatom frustule; 2 stands for considering the two sides of the frustule assuming a similar distribution of the attached bacteria on both sides; n is the number of attached bacteria detected on one frustule side; *c* is a coefficient calculated from the ratio of the nucleus area covered by the DAPI signal of the diatom and the visible area of diatom frustule. The mean value of this coefficient for *T. rotula* cells in this study is 0.12 (range from 0.04 to 0.32). It was determined from measurements of diatom cells collected on days 1, 7, 15 and 21 (60 cells at each time point; Supporting Information Description S1). The numbers of target bacteria detected by FITC signals were calculated similarly to that of total colonizers. However, because FITC signals of bacterial cells were not masked by signals of the diatom nucleus like in the DAPI channel (Supporting Information Fig. S2), the coefficient *c* was considered as 0 and the formula N = 2 × n was applied to calculate the number of target bacteria on their host diatom.

### Assessing of diatom-attached bacteria using suspended samples

Fixed culture samples of the consortium treatment stored in 2 ml safe-lock tubes were used to analyze the attached bacteria by a CARD-FISH procedure modified from current protocols (Pernthaler et al. 2002, Bakenhus et al. 2017) as follows and described in Table 2. Centrifugation steps were applied to isolate the diatoms with the attached bacterial cells from the culture suspension from which subsamples were withdrawn by pipette. To limit the damage of the diatom cells as well as the loss of the bacteria attached to the diatom, a series of time and speed parameters of centrifugation were pretested to obtain the suitable values of 3 - 5 min and 3000 g, respectively. Before hybridization, samples were dehydrated and autofluorescence of chlorophyll reduced with increasing concentrations of ethanol (70%, 80% and 96% [v/v]). Probes used in this experiment were CF319a and GAM42a (Table 1). Probe concentrations of 0.5, 1.0, 2.0 and 5 ng µl^-1^ were tested and evaluated to choose the most effective concentrations for the experiment. The final concentration of 5 ng µl^-1^ in hybridization buffer (35% FA) was found as the optimal value.

**Table 2.**
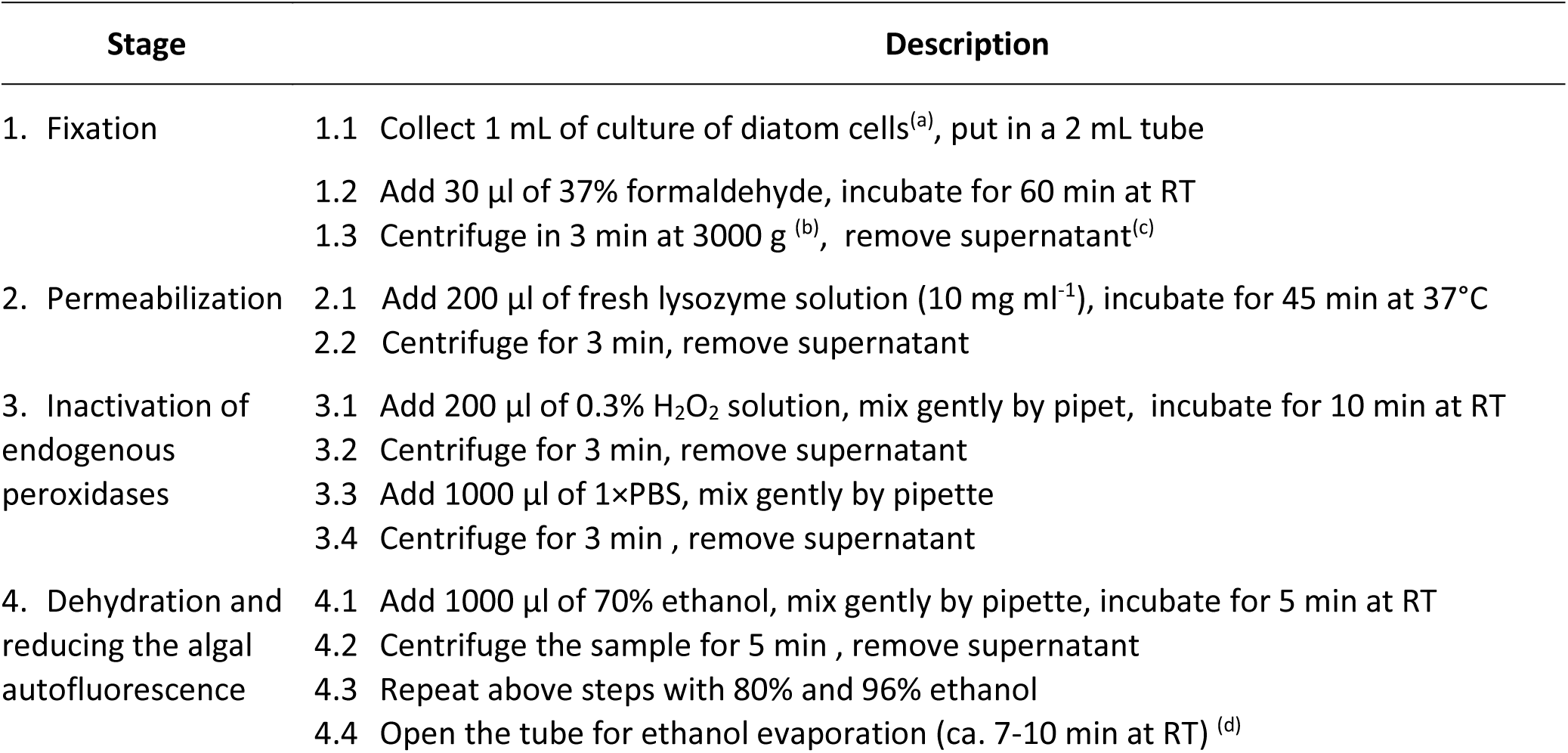
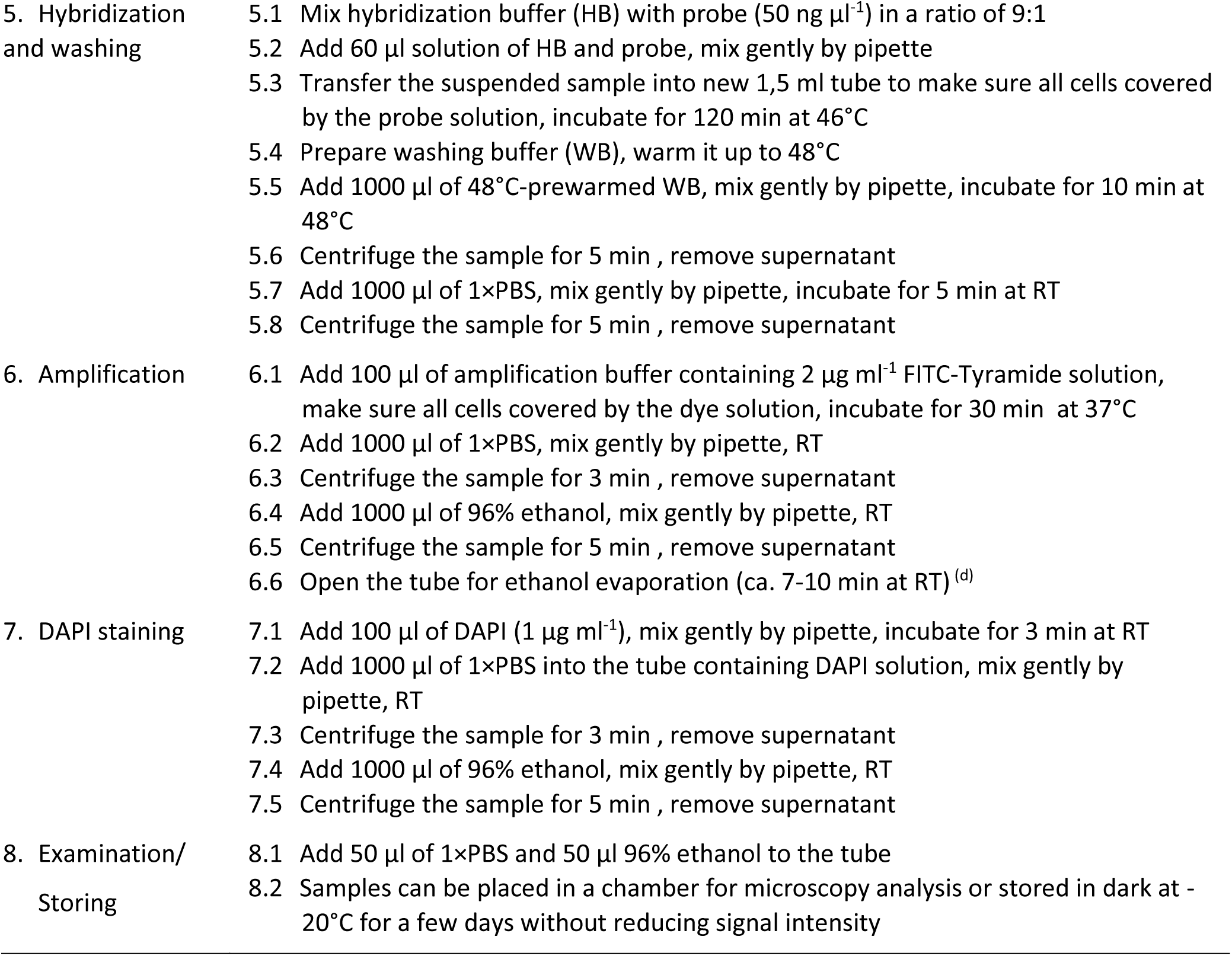
Improved CARD-FISH protocol to enumerate attached bacteria on *T. rotula* using suspended samples.

| Stage | Description |
| --- | --- |
| 1. Fixation | 1.1 Collect 1 mL of culture of diatom cells<sup>(a)</sup>, put in a 2 mL tube<br>1.2 Add 30 $\mu\text{l}$ of 37% formaldehyde, incubate for 60 min at RT<br>1.3 Centrifuge in 3 min at 3000 g<sup>(b)</sup>, remove supernatant<sup>(c)</sup> |
| 2. Permeabilization | 2.1 Add 200 $\mu\text{l}$ of fresh lysozyme solution (10 mg $\text{mL}^{-1}$), incubate for 45 min at 37°C<br>2.2 Centrifuge for 3 min, remove supernatant |
| 3. Inactivation of endogenous peroxidases | 3.1 Add 200 $\mu\text{l}$ of 0.3% $\text{H}_2\text{O}_2$ solution, mix gently by pipet, incubate for 10 min at RT<br>3.2 Centrifuge for 3 min, remove supernatant<br>3.3 Add 1000 $\mu\text{l}$ of 1×PBS, mix gently by pipette<br>3.4 Centrifuge for 3 min, remove supernatant |
| 4. Dehydration and reducing the algal autofluorescence | 4.1 Add 1000 $\mu\text{l}$ of 70% ethanol, mix gently by pipette, incubate for 5 min at RT<br>4.2 Centrifuge the sample for 5 min, remove supernatant<br>4.3 Repeat above steps with 80% and 96% ethanol<br>4.4 Open the tube for ethanol evaporation (ca. 7-10 min at RT)<sup>(d)</sup> |

|  |  |  |
| --- | --- | --- |
| 5. Hybridization and washing | 5.1 | Mix hybridization buffer (HB) with probe (50 ng $\mu\text{l}^{-1}$) in a ratio of 9:1 |
| | 5.2 | Add 60 $\mu\text{l}$ solution of HB and probe, mix gently by pipette |
|  | 5.3 | Transfer the suspended sample into new 1,5 ml tube to make sure all cells covered by the probe solution, incubate for 120 min at 46°C |
|  | 5.4 | Prepare washing buffer (WB), warm it up to 48°C |
| | 5.5 | Add 1000 $\mu\text{l}$ of 48°C-prewarmed WB, mix gently by pipette, incubate for 10 min at 48°C |
|  | 5.6 | Centrifuge the sample for 5 min , remove supernatant |
| | 5.7 | Add 1000 $\mu\text{l}$ of 1×PBS, mix gently by pipette, incubate for 5 min at RT |
|  | 5.8 | Centrifuge the sample for 5 min , remove supernatant |
| 6. Amplification | 6.1 | Add 100 $\mu\text{l}$ of amplification buffer containing 2 $\mu\text{g ml}^{-1}$ FITC-Tyramide solution, make sure all cells covered by the dye solution, incubate for 30 min at 37°C |
| | 6.2 | Add 1000 $\mu\text{l}$ of 1×PBS, mix gently by pipette, RT |
|  | 6.3 | Centrifuge the sample for 3 min , remove supernatant |
| | 6.4 | Add 1000 $\mu\text{l}$ of 96% ethanol, mix gently by pipette, RT |
|  | 6.5 | Centrifuge the sample for 5 min , remove supernatant |
|  | 6.6 | Open the tube for ethanol evaporation (ca. 7-10 min at RT) <sup>(d)</sup> |
| 7. DAPI staining | 7.1 | Add 100 $\mu\text{l}$ of DAPI (1 $\mu\text{g ml}^{-1}$), mix gently by pipette, incubate for 3 min at RT |
| | 7.2 | Add 1000 $\mu\text{l}$ of 1×PBS into the tube containing DAPI solution, mix gently by pipette, RT |
|  | 7.3 | Centrifuge the sample for 3 min , remove supernatant |
| | 7.4 | Add 1000 $\mu\text{l}$ of 96% ethanol, mix gently by pipette, RT |
|  | 7.5 | Centrifuge the sample for 5 min , remove supernatant |
| 8. Examination/<br>Storing | 8.1 | Add 50 $\mu\text{l}$ of 1×PBS and 50 $\mu\text{l}$ 96% ethanol to the tube |
|  | 8.2 | Samples can be placed in a chamber for microscopy analysis or stored in dark at -20°C for a few days without reducing signal intensity |

Samples within tubes after CARD-FISH and DAPI-staining were placed on microscope slides with reaction wells (Paul Marienfeld, Lauda-Koenigshofen, Germany) and examined using the microscopic system as described above (Carl Zeiss Microscope AxioImager.Z2m, objective 63x NA 1.46, black and white AxioCam MRm camera, and AxionVision program). In addition, to evaluate the cell loss of diatom-attached bacteria, *T. rotula* cells and epibionts after CARD-FISH and DAPI-staining were filtered onto white polycarbonate filters (5 µm, Isopore). The quantification of attached bacteria was done as described for the filter sections. The comparison of the numbers of attached bacteria by the filtered and centrifuged samples yielded the loss of attached cells by centrifugation.

### Statistical analysis

Statistical analyses to evaluate the significance of the results were performed using one way analysis of variance (ANOVA). Pair-wise comparison procedures were conducted following the significant ANOVA by a Tukey test. Significant differences were considered at *p*≤0.05. The statistical analysis software SigmaPlot 12.0 (Systat Software, San Jose, USA) was used for test calculations.

## Results

We set up experiments with an axenic strain of *T. rotula* isolated from the North Sea (Tran et al. 2022) which was inoculated either with a consortium of five bacterial strains or with a natural bacterial community collected from the German Wadden Sea and incubated at 15°C in a 12:12 h light:dark cycle for 21 days. The consortium included *Gramella forsetii* (*Bacteroidetes*), *Glaciecola* sp. (*Gammaproteobacterium*), *Colwellia* sp. M166 (*Gammaproteobacterium*), *Roseovarius* sp. M141 (*Roseobacter*) and *Pseudophaeobacter* sp. (*Roseobacter*). Samples for CARD-FISH analyses were withdrawn at days 10 and 21 and processed as described below.

### Assessment of bacterial colonizers on *T. rotula* by CARD-FISH using filtered samples

CARD-FISH analysis of filtered samples containing bacteria attached to diatom cells revealed clear signals of both DAPI-stained and hybridized bacteria (Fig. 1, Supporting Information Fig. S3). There was no difference in the intensity of FISH signals obtained by probe concentrations of 0.5 or 1 ng µl^-1^. Although non-specific chlorophyll fluorescence of a few *T. rotula* cells interfered with DAPI-stained as well as FISH-signals of the epibionts (Supporting Information Fig. S4), attached cells were clearly visible and easily counted. In each field of view (FOV), the stacked image after the multilayer imaging process made it possible to count most of the bacterial cells attaching to frustules of *T. rotula* (Fig. 1A). In the DAPI channel, some bacterial cells were not visible due to the nuclear signals of the diatom, but these signals were mostly countable in the FITC channel (Figs. D2 and E2).

**Figure 1.**
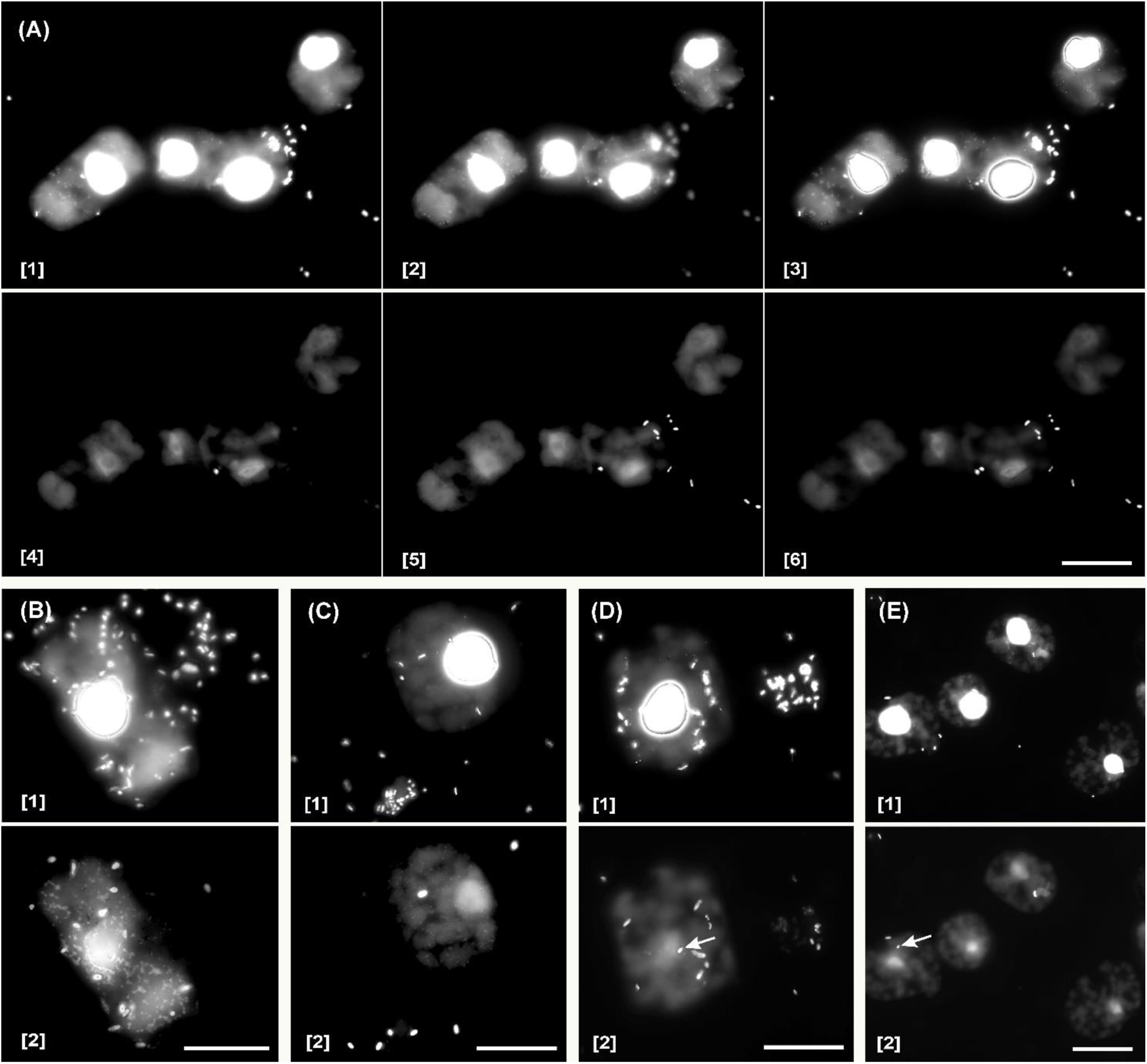
Micrographs of DAPI-stained and hybridized cells of the natural bacterial community and species consortium attached to *T. rotula* detected by the CARD-FISH assay using filtered samples. (A) Cells of the natural bacterial community sampled at day 10 and hybridized with probe CF319a; (A3) the stacked DAPI-image of Figs. A1 and A2; (A6) the stacked FISH-image of Figs. A4 and A5. (B) Cells of the natural bacterial community samples at day 21 and hybridized with probe ROS536. (C-E) Cells of the consortium hybridized with probes CF319a (C), GAM42a (D) and ROS536 (E). (B1, C1, D1, E1) stacked DAPI-images. (B2, C2, D2, E2) stacked FISH-images. The arrows in Figs. D2 and E2 indicate attached bacteria which were not visible in the corresponding DAPI images due to DAPI-signals of the diatom nuclei. Raw data from different focal planes of merging images B-E are showed in the Supporting Information Fig. S3. Scale bar = 20 µm.

At abundances of attached bacteria below 10 cells frustule^-1^ (Fig. 1C, 1E), they could be enumerated directly without stacked and merged micrographs. In contrast, at high abundances of attached bacteria (e.g., Fig. 1A, 1B, 1D) stacking of 2-5 micrographs from different focal planes in one FOV and generating a composite micrograph by the Picolay software (www.picolay.de) was necessary to assess numbers of attached bacteria. Only by this approach all attached bacteria located on one half of the algal frustule became clearly visible and thus countable (e.g., Fig. 1A, Supporting Information Fig. S3).

Applying this counting approach it was evident that numbers of bacterial colonizers on *T. rotula* frustules in both treatments increased significantly with incubation time (Figs. 2A and 2B). At day 10, the number of attached DAPI-stained cells in the natural bacterial community was twice as high as that in the species consortium (12.5 ± 3.3 compared to 6.0 ± 1.9 cells frustule^-1^) and this difference increased to more than 10-fold at day 21 (171.0 ± 22.7 compared to 14.6 ± 1.9 cells frustule^-1^). In samples of the natural bacterial community from the latter day many *T. rotula* frustules which were transparent or broken exhibited high abundances of attached bacteria (Supporting Information Fig. S5). In line with total bacterial numbers the cell numbers of distinct bacterial strains detected by specific probes increased also over time with 2 to 20 times higher values at day 21 compared to those at day 10 (Figs. 2A and 2B, *p*<0.001). In the consortium treatment, attached cells of *Gammaproteobacteria* were numerically dominant, showing higher abundances than those of the two other bacterial groups, *Bacteroidetes* and *Roseobacter*, (*p*<0.05) at both sampling points, accounting for 3.9 ± 0.4 cells frustule^-1^ (57.6 ± 6.5%) and 6.4 ± 0.8 cells frustule^-1^ (42.7 ± 3.9%) at days 10 and 21, respectively. By contrast, *Bacteroidetes* showed lowest abundance, contributing only 0.3 ± 0.1 cells frustule^-1^ (4.7 ± 1.5%) at day 10 and 2.9 ± 0.8 cells frustule^-1^ (17.9 ± 4.1%) at day 21. In the treatment with the natural community, there were no significant differences in the abundances of the three bacterial groups at day 10 (*p*=0.156), but at day 21 *Roseobacter* members were the most dominant colonizers (61.0 ± 12.5 cells frustule^-1^; 33.2 ± 1.5%) while *Gammaproteobacteria* showed the lowest numbers (11.6 ± 3.3 cells frustule^-1^; 6.6 ± 1.6%; *p*<0.05).

**Figure 2.**
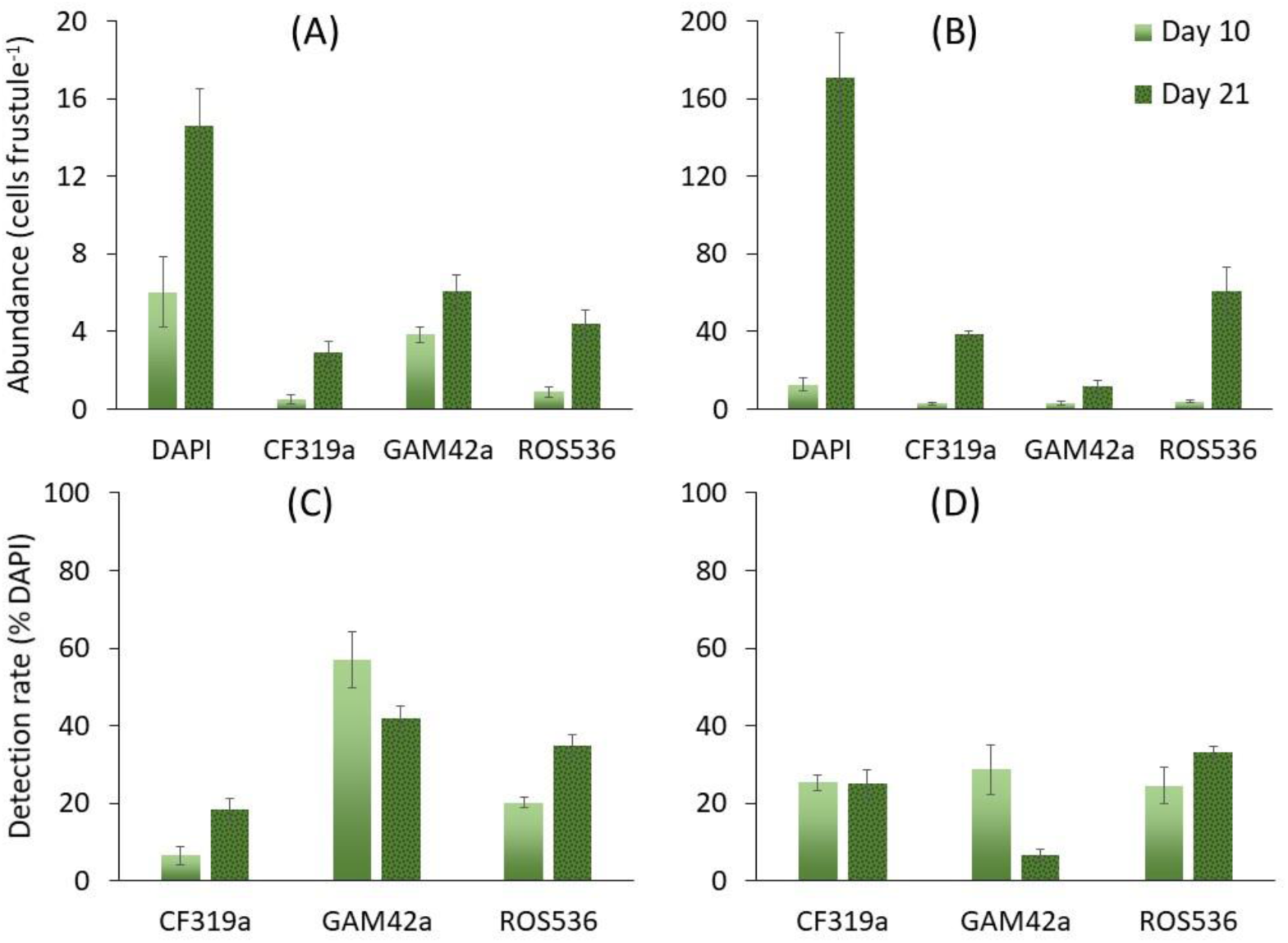
Abundances of bacteria attached to *T. rotula* at days 10 and 21 detected by CARD-FISH with probes CF319a, GAM42a and ROS536 using filtered samples. (A, B) Total DAPI cell counts and absolute abundances (cells frustule^-1^) of target bacteria in the consortium (A) and treatment with a natural community (B). Note the different scale in (B). (C, D) Relative abundances (% DAPI) of target bacteria in the consortium (C) and treatment with a natural community (D).

The proportions of attached bacteria (% DAPI) were different in both treatments and did not follow the trend of the absolute numbers (Figs. 2C and 2D). The proportions of *Bacteroidetes* and *Roseobacter* members increased with incubation time, in line with the absolute numbers. In contrast, the fraction of *Gammaproteobacteria* decreased sharply at day 21 in both treatments, despite increasing absolute numbers. Total relative abundance of hybridized bacteria detected by the three probes (CF319a, GAM42a and ROS536) accounted for 83.8% of attached DAPI-stained cells at day 10 and 95.3% at day 21 in the consortium treatment, and 78.5% at day 10 and 64.8% at day 21 in the treatment with the natural community.

### Examination of attached bacteria on *T. rotula* by CARD-FISH using suspended samples

The CARD-FISH approach using cells in suspension with subsequent concentration by centrifugation with probe concentrations of 0.5 or 1.0 ng µl^-1^ showed very weak signals in the bacteria of all samples (Supporting Information Fig. S6). In contrast, the results of probes CF319a and GAM42a with the concentration of 2.0 or 5.0 ng µl^-1^ showed conspicuous signals (Fig. 3; Supporting information Fig. S7). Similarly to filtered samples, some *T. rotula* cells showed bright nonspecific fluorescence that could obscure or interfere with positive bacterial signals in both DAPI and FITC channels (Supporting Information Fig. S4). Our comparative observations of the filtered and centrifuged samples revealed that centrifugation and resuspension steps during the hybridization procedure did not damage the frustules of *T. rotula*, but removed the algal threads and washed away free-living bacterial cells. Indeed, bacterial as well as non-specific fluorescence signals surrounding the host frustules were not found after CARD-FISH as observed in the filtered samples on the membrane sections. At the centrifugation parameters we used, diatom cell clumps were not observed after sample processing. Because this study did not examine the host cell abundance, the quantification of the diatom was not carried out. The background of the images contained less noise than those obtained by the filtered samples after CARD-FISH processing (e.g., Figs. 1B-1D).

**Figure 3.**
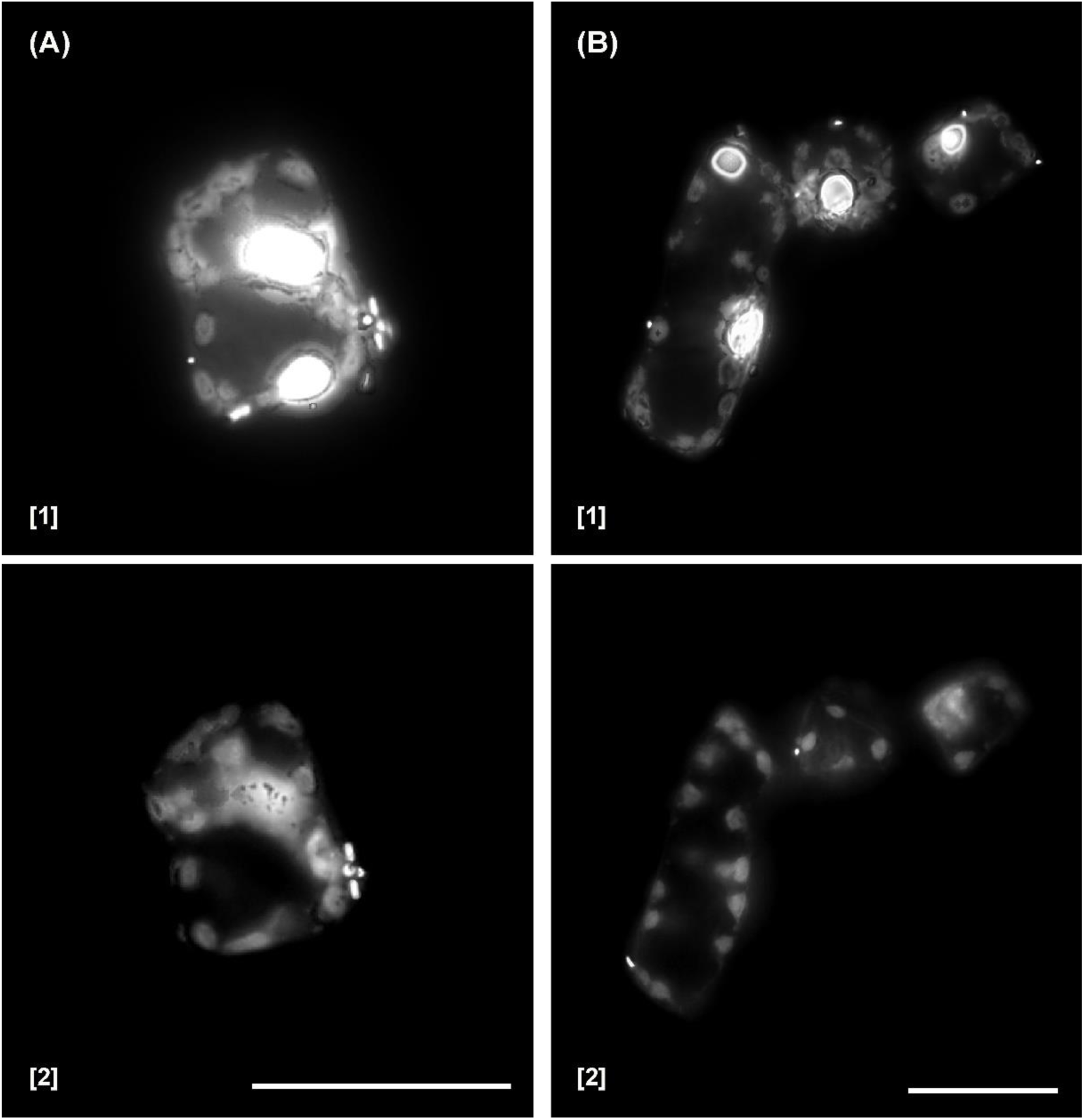
Micrographs of attached DAPI-stained and hybridized bacteria on *T. rotula* frustules detected by CARD-FISH with probe CF319a (A) and GAM42a (B) using suspended samples of species consortium treatment (day 21). (A1 and B1) Stacked DAPI-images. (A2 and B2) Stacked FISH-images. Raw data from different focal planes of merging images A and B are showed in the Supporting Information Fig. S7. Scale bar = 20 µm.

To assess the possible loss of attached bacteria caused by CARD-FISH processing of the samples in suspension, we quantified the attached bacteria on diatom frustules in a similar way as that applied on filtered samples. The absolute abundance of attached DAPI-stained cells, *Bacteroidetes* and *Gammaproteobacteria* detected in suspended samples collected at day 21 was 14.7 ± 2.6, 3.0 ± 0.3 and 6.5 ± 3.9 cells frustule^-1^, respectively. These numbers were not significantly different from the corresponding numbers obtained from filtered samples after CARD-FISH processing (*p*>0.05).

## Discussion

The results of this study indicate that this CARD-FISH procedure using filtered or suspended and centrifuged samples and reducing chlorophyll fluorescence of the diatom cells is well suitable for the identification and quantification of bacteria attached to diatoms such as *T. rotula*. The technique with tomography-like stacked micrographs allowed merging images of several focal planes of captured bacteria attached to the diatom frustules into a composite image with all bacteria of one side of the diatom cell well in focus. The low chlorophyll fluorescence by an extra ethanol treatment prior to the hybridization procedure made it possible to enumerate total attached bacterial cells after DAPI staining and to apply probes fluorescing in the visible light for inspecting and enumerating bacteria on the diatom frustules. Previous methods circumvented the problem of high chlorophyll autofluorescence by applying probes emitting fluorescence in the infrared light like Cy5 (Wada et al. 2016) or completely remove cellular pigments (Castillo et al. 2020). Other studies assessing bacterial colonization of macroalgae (Tujula et al. 2006, Brunet et al. 2021) and a picoplankton alga (Castillo et al. 2020) by CARD-FISH also successfully bleached chlorophyll fluorescence by alcohol treatment but the colonization patterns on the macroalgal surfaces are more like a biofilm structure than on single microalgal cells. For examining bacteria colonizing diatoms or other microalgal cells it is a great advantage when the host cell is visible simultaneously with the targeted attached bacteria labelled with a fluorescent probe in the visible light and DAPI counterstaining which, with a probe with infrared fluorescence, is only possible by image analytical processing, or by eliminating pigments.

In the *T. rotula* culture with the species consortium, the bacterial numbers targeted with the three probes CF319a, GAM42a and ROS536 together reached 83.8% and 95.3% of the total DAPI-stained cells on the diatoms at days 10 and 21, respectively, and in that with the natural bacterial community 64.8% and 78.5%, respectively. This emphasizes that our CARD-FISH method yields high and reliable numbers by facilitating the hybridization procedure. Theoretically, total numbers of hybridized bacteria in the consortium treatment detected by probes CF319a, GAM42a and ROS536 should be equivalent to the total number of DAPI-stained cells. However, a small portion of DAPI signals detected in this treatment was not recognized as FISH signals after the CARD-FISH approach. This is presumably because genetic material of the diatom leaked out, spread onto the frustules and exhibited similar signals as bacteria, leading to higher numbers of DAPI-stained structures than the actual number of attached bacteria. Further, some fluorescently labelled particles, accounting in total for <10% of total particles, emerged as little spots on the frustules during the imaging process, which could obviously not be distinguished from the bacterial DAPI signals, were considered as epibionts. We do not know how and from which material these little spots were formed. On the other hand, non-specific fluorescence appearing in the FITC channel may have been so strong that it obscured some of positive FISH signals, leading to lower cell numbers of the target bacteria. Similar constraints have been reported from previous studies (Castillo et al. 2020, Piwosz et al. 2021). In one study which applied the CARD-FISH method to visualize the food vacuole content of heterotrophic flagellates, weak or lack of signals, high background fluorescence and non-specific probe binding were reported (Piwosz et al. 2021). These authors mention as possible reasons causing these problems poorly designed probe, low fluorescence activity or loose binding of the probe and low ribosome content of the bacterial target groups. Another study (Castillo et al. 2020) reduced the high background chlorophyll fluorescence by immersing the sample or filter in 96% ethanol and then in pure methanol. Another way to overcome the intense autofluorescence background noise was to create strong CARD-FISH signals by high concentrations of tyramide in benthic cyanobacterial mats (Abed et al. 2002).

The fraction of the bacteria in the natural community detected with the three probes was lower than in the species consortium. This may reflect that, in addition to the shortcomings mentioned above for the consortium, other prokaryotes not targeted by the probes applied, such as *Archaea*, *Verrucomicrobiales* and *Planctomycetes*, may have been also members of the natural prokaryotic diatom-associated prokaryotic community (Herfort et al. 2007, Jarrell et al. 2013, Bižić-Ionescu et al. 2015).

In the filtered diatom samples, there was no significant difference of the background signal applying different probe concentrations (0.5 to 5.0 ng µl^-1^). This observation is in contrast to a previous report that high probe concentration can cause background fluorescence obscuring enumeration of bacterial cells (Pernthaler et al. 2002). In our study, we examined bacteria attached to diatom frustules such that non-specific fluorescence caused by other factors than background fluorescence, like of chlorophyll, appeared to be more important than probe concentration.

It has been shown by CLSM that diatom cells are surrounded by threads or setae consisting of glycoconjugate structures which can be visualized by fluorescent lectins (Herth and Barthlott 1979, Kaczmarska et al. 2005, Bennke et al. 2013, Tran et al. 2022). These threads are loosely attached to the diatom cells and also colonized by bacteria and can be easily detached from the cells. As bacteria attached to the threads are not clear-cut attached to the diatom frustules we removed the potentially present threads by washing of the filtered samples. Further, bacterial cells were counted only if they were identified as directly attached to the diatom frustule.

The well-focused stacked images with the bright and identifiable bacterial cells attached to the *T. rotula* frustules clearly showed the significant increase of attached bacterial numbers in the species consortium and the natural community from day 10 to 21. This indicates that our CARD-FISH protocol is very suitable to examine the colonization pattern of bacterial communities on host diatom cells in the course of a growing co-culture and presumably also during a natural diatom bloom and possibly also on diatom-dominated biofilms. In addition, the enumeration of epibionts which revealed low abundance of *Bacteroidetes* (0.3 ± 0.1 cells frustule^-1^; 4.7 ± 1.5%) in the consortium treatment and *Gammaproteobacteria* in the natural treatment (11.6 ± 3.3 cells frustule^-1^; 6.6 ± 1.6%) further indicate that the approach is useful for the examination of colonizer communities with members of low abundances.

The fluorescence of DAPI-stained algal nuclei might mask signals of attached bacteria leading to underestimating their number. To take into account this potential bias, counted numbers of DAPI-stained bacteria were corrected by a factor considering the area of the DAPI-stained nucleus. Hence, based on the determination of this area, a coefficient *c* and a formula was used to calculate absolute numbers of bacteria attached to the diatom frustules. Further, to consider both sides of the host cell numbers were doubled. This procedure to calculate absolute numbers of bacteria attached to diatom cells differs from previous publications, which reported results as determined from one side of the host cell (Vaqué et al. 1989, 1990, Arandia-Gorostidi et al. 2022). We propose to use this conversion for future investigations with CARD-FISH and DAPI counterstaining not to underestimate the total number of bacteria attached to diatoms.

In the CARD-FISH analysis performed with diatom cells in suspension, low probe concentrations, i.e. 0.5 to 1.0 ng ml^-1^, did not yield fluorescence strong enough for a reliable signal assessment, in contrast to the results obtained with the filtered samples. This may be because prior to hybridization, samples in the tubes were not completely dried leading to a possible further dilution of the probe concentration in the hybridization buffer. Another possibility is that the dehydration of cells with ethanol may have been incomplete, leading to remains of water in the target bacterial cell, thereby reducing the permeability of the probe solution. Nevertheless, high probe concentrations (2 or 5 ng µl^-1^) yielded reliable results and the concentration of 5 ng µl^-1^ was considered as the optimal value. This is in agreement with previous studies applying FISH (Böckelmann et al. 2002, Batani et al. 2019) or CARD-FISH analysis (Biegala et al. 2003, Manti et al. 2011) with unfiltered samples.

## Conclusion

Our results showed in the diatom co-culture both with the species consortium and the natural bacterial community similar numbers of bacteria attached to cells of *T. rotula* by both concentration methods we used, filtration and centrifugation of suspended samples. With filtered samples, numbers of attached bacteria were examined at low numbers without any further manipulation after CARD-FISH and DAPI staining in one focal plane by epifluorescence microscopy and with high numbers on a composite image after stacking of multiple micrographs. In this setting, free-living bacteria and those loosely attached to the diatom and its surface structures such as setae and threads can be examined as well. In contrast, in the centrifuged samples the latter bacteria were not assessable. However, the clear DAPI and CARD-FISH signals of the attached bacteria on the tomographic-liked 3-dimensional images of the stacked micrographs allow a highly resolved examination of different populations of bacteria attached to the frustules of *T. rotula* and presumably to other diatoms and possibly also other microalgae with firm surface structure. Hence our improvements of the CARD-FISH method are useful and presumably superior to previous methods to assess the colonization of diatoms and presumably other microalgae with firm cell structures in co-cultures with bacteria and in the course of natural phytoplankton blooms.

## Acknowledgments

The authors greatly appreciate Rolf Weinert for maintaining and providing axenic culture of the diatom *T. rotula* and Mathias Wolterink for preparing bacterial strains for the co-cultures. We thank Bernhard Fuchs for his valuable comments on an earlier version of this manuscript. This study was supported by Deutsche Forschungsgemeinschaft (DFG) within the Transregional Collaborative Research Centre “*Roseobacter*” (TRR51) and Deutscher Akademischer Austauschdienst (DAAD).

## Author contributions

TQD and MS designed the study. TQD carried out the experiments, the microscopic studies, analyzed the data and drafted the manuscript. MS contributed to data analysis and writing the.

## Supporting Information

**Description S1.** Determination of c value for calculation of DAPI-stained cell number.

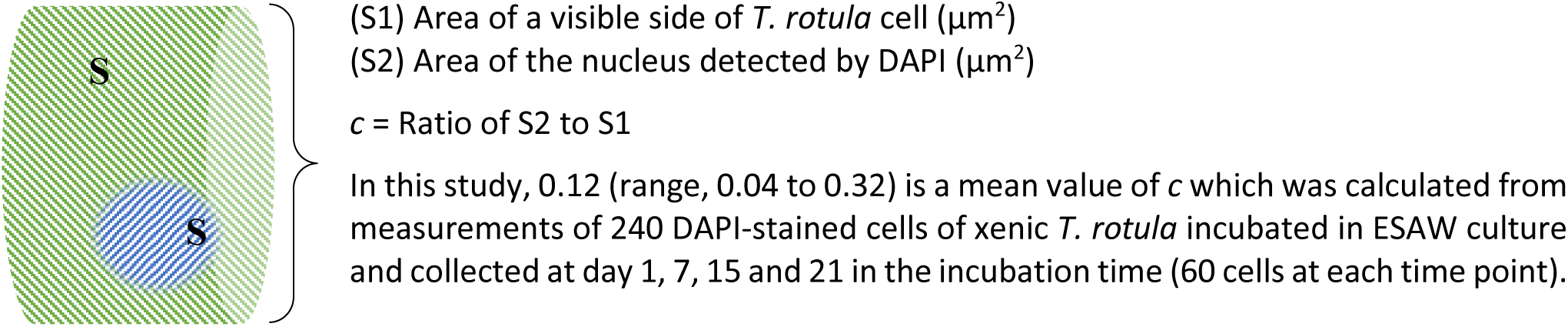

*Example:*

Calculate DAPI-stained cell number (N) on 2 sides of a frustule when emerged DAPI signals is 8 (n = 8)

=> Calculated number of DAPI-stained cells on 2 sides of a frustule N = 2 × (n + (*c* × n)) = 2 × (8 + 8 × 0.12) = 17.92 cells frustule^-1^

**Figure S1.**
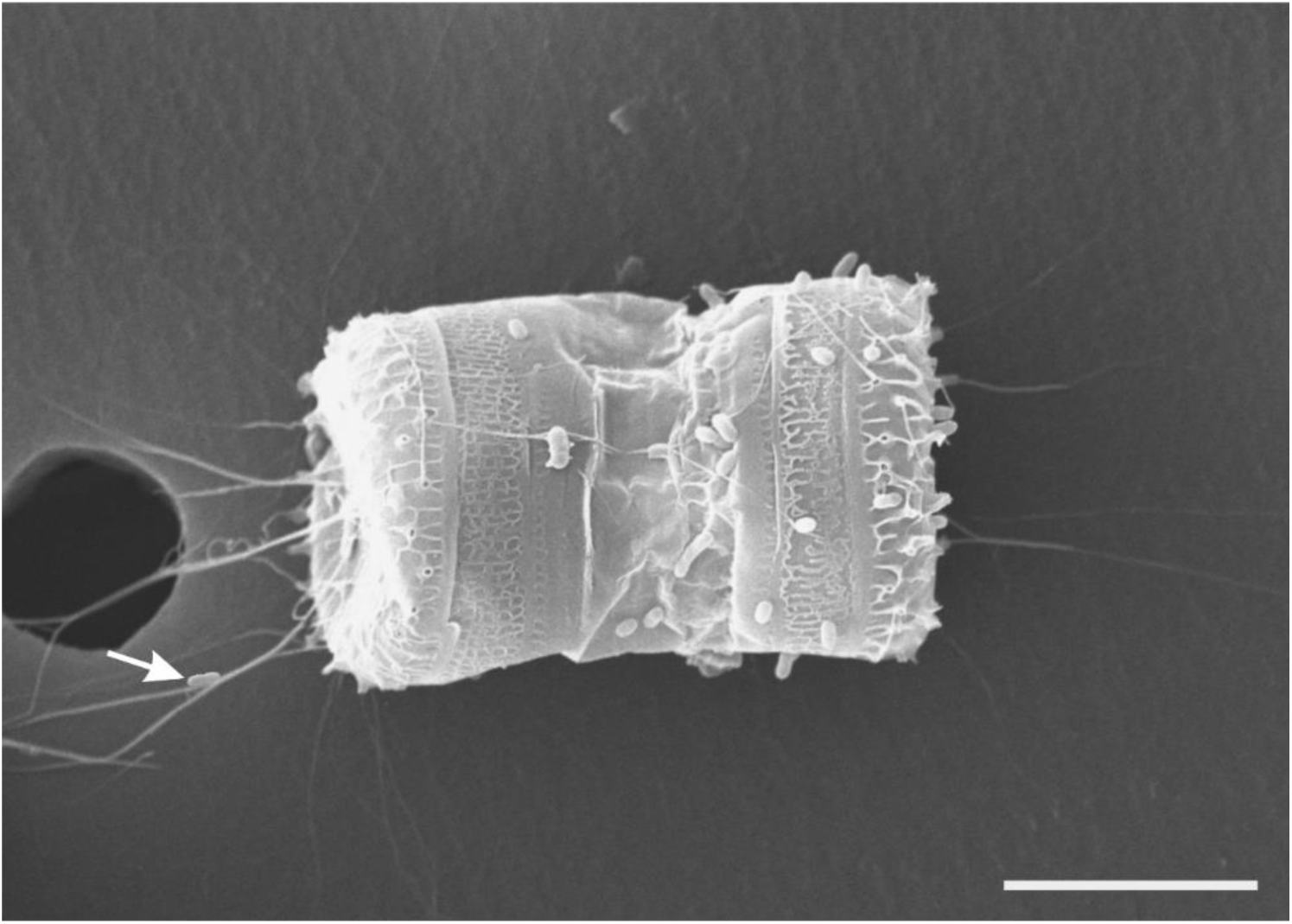
Scanning electron micrograph of a xenic *T. rotula* cell showed attached bacteria on the frustule as well as the threads of a host diatom cell. The arrow indicates attached bacteria on the threads. Scale bar = 6 µm.

**Figure S2.**
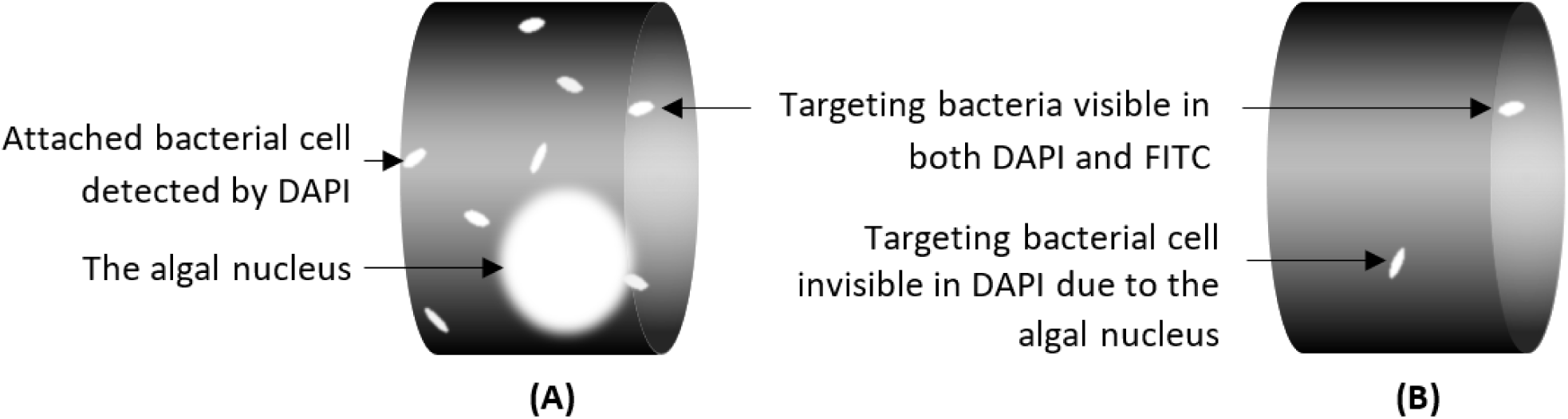
The scheme of a host *T. rotula* frustule and attached bacteria under a fluorescent microscope. (A) DAPI signals of the algal nucleus and total cell counts, (B) FITC signals of targeting attached bacteria.

**Figure S3.**
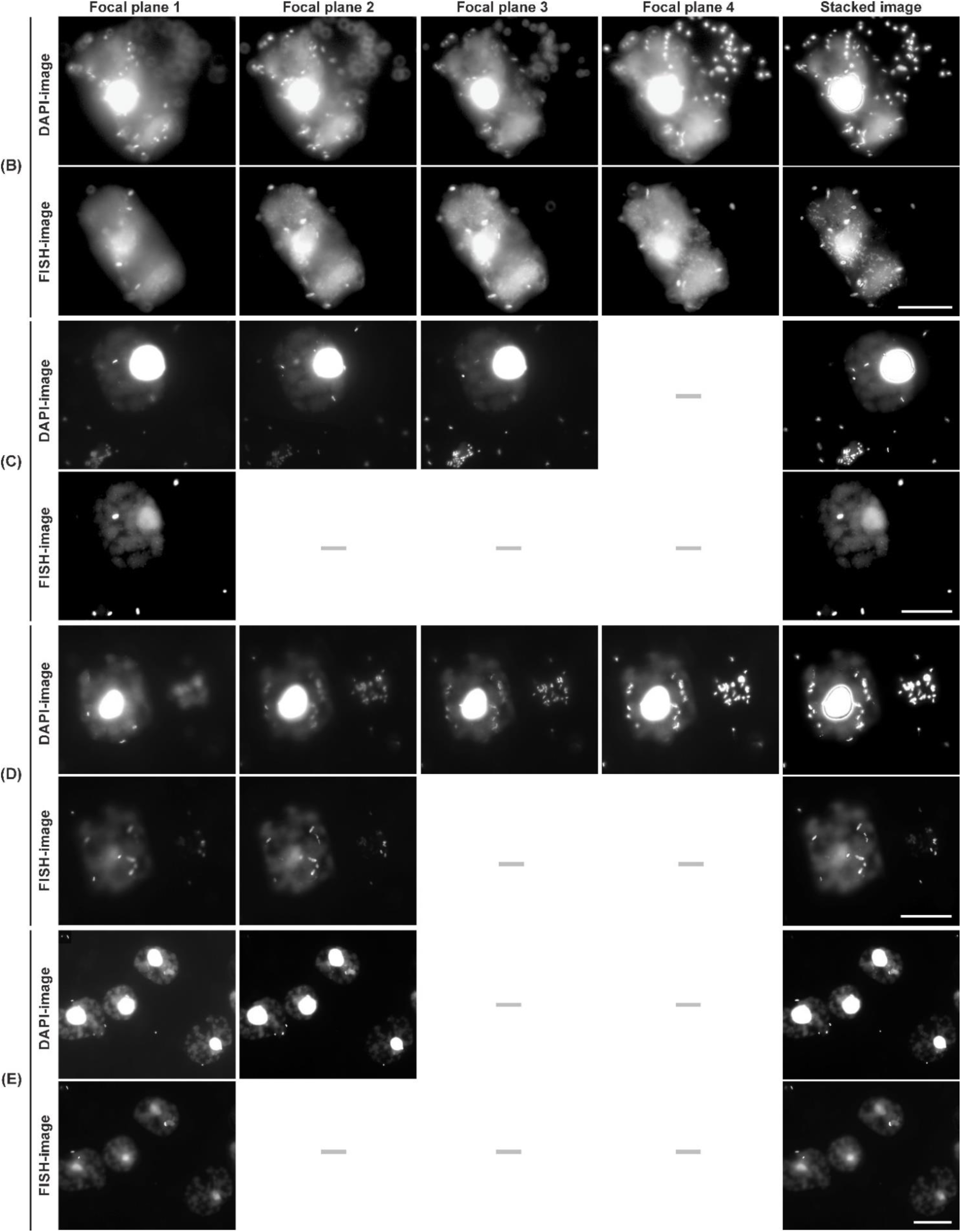
Raw images from different focal planes of Figures 1B-1E (stacked images). The sign “–“ means that the image was not taken at the corresponding focal plane. Scale bar = 20 µm.

**Figure S4.**
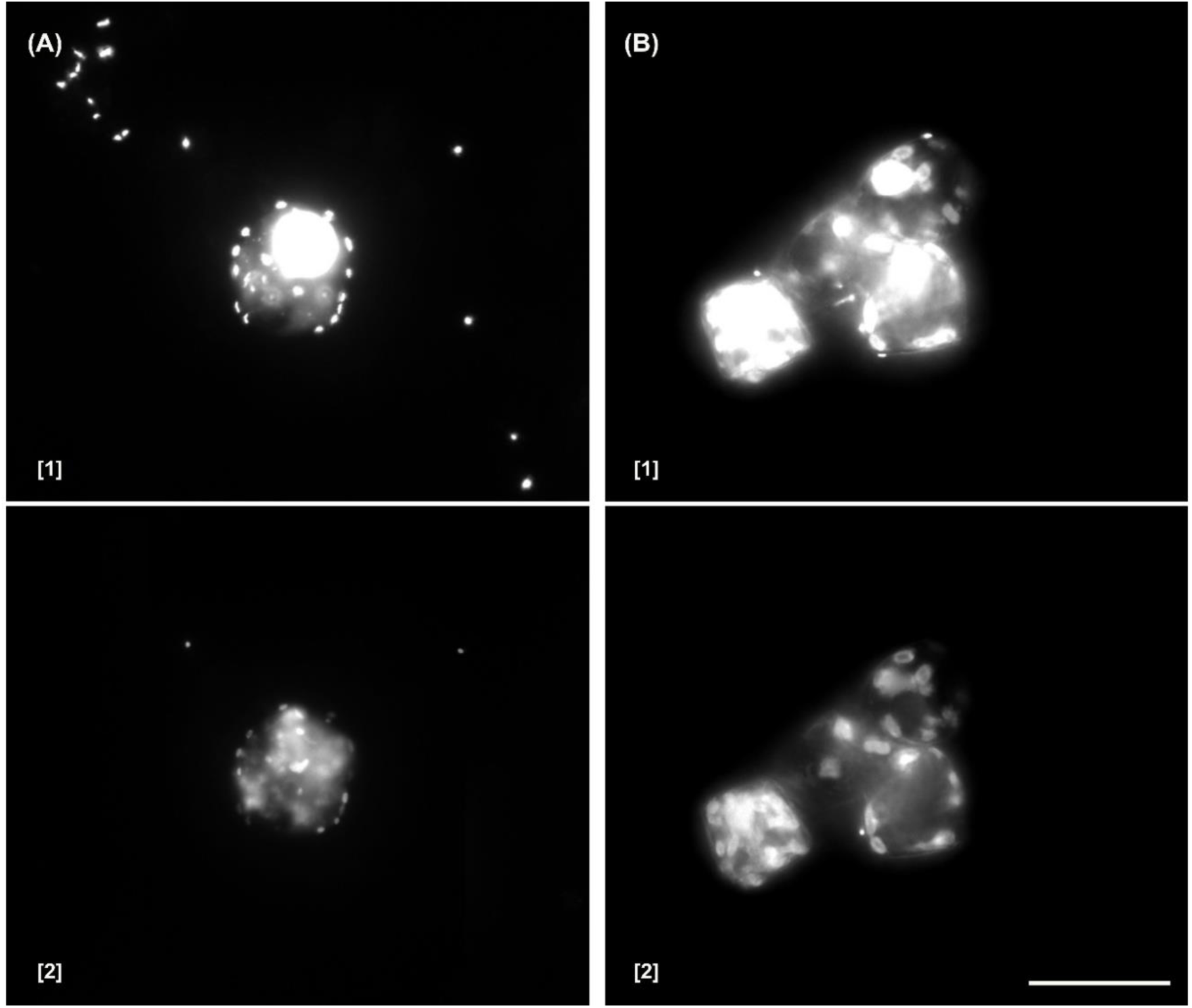
Attached bacteria on a diatom cells after CARD-FISH using filter membrane (A) and tube (B) samples with probe GAM42a. (A) DAPI image showing clear signals of DAPI-stained bacteria and the diatom nucleus (A1), FITC image showing the algal autofluorescence which interfered with FISH-signals (A2). (B) DAPI image showing unspecific fluorescent signals which interfered with DAPI-signals (B1), FITC image showing clear signals of *Gammaproteobacteria* (B2). Scale bar = 20 µm.

**Figure S5.**
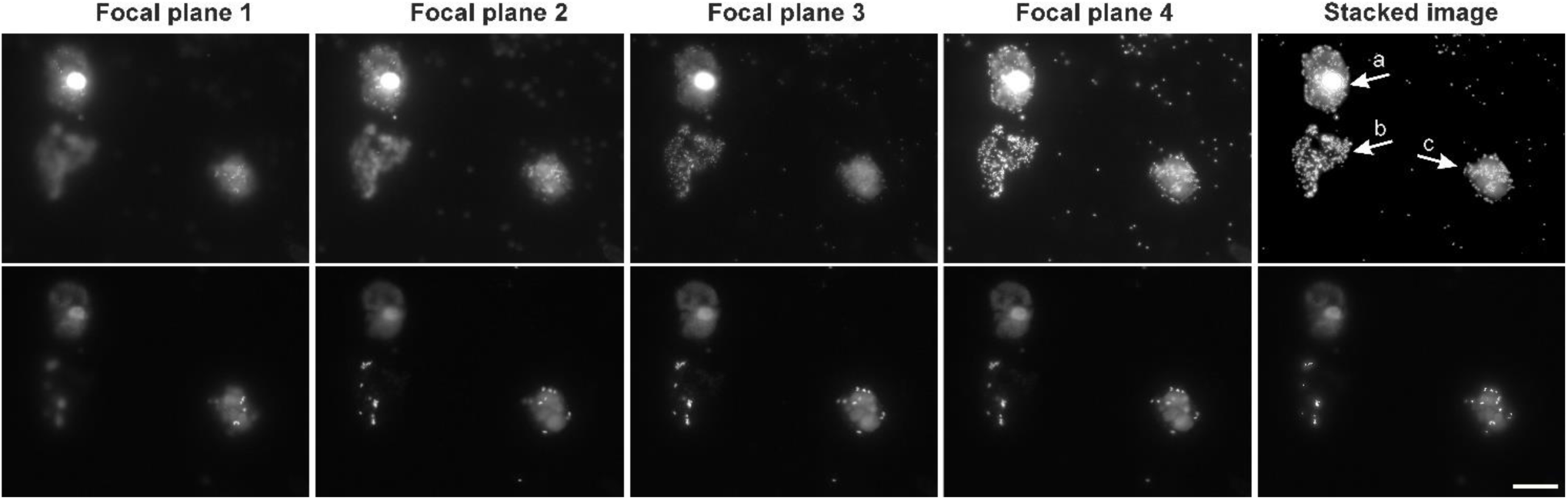
High density of attached bacterial cells on intact (a), broken (b) and transparent (c) frustules of *T. rotula* in natural treatment after CARD-FISH with probe ROS536 using a membrane filtered sample. DAPI images on the top and FISH images on the bottom. Scale bar = 20 µm.

**Figure S6.**
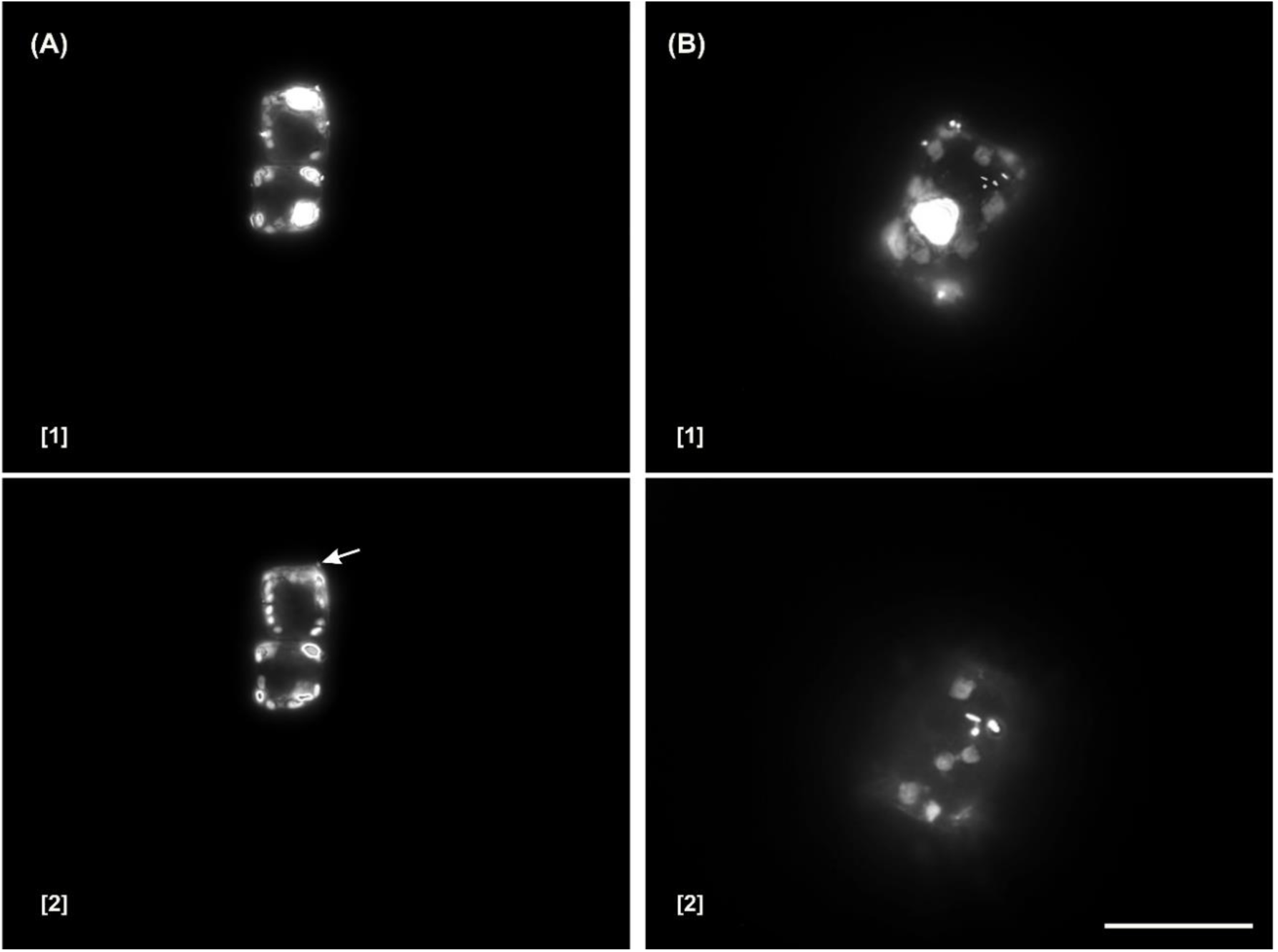
Attached *Gammaproteobacteria* cells on frustules of *T. rotula* after CARD-FISH using probe GAM42a with the concentration of 0.5 ng µL^-1^ (A2) and 5 ng µL^-1^ (B2). (A1 and B1) The DAPI stacked images. (A2 and B2) The FITC stacked images. The arrow indicates weak FISH-signals. Scale bar = 20 µm.

**Figure S7.**
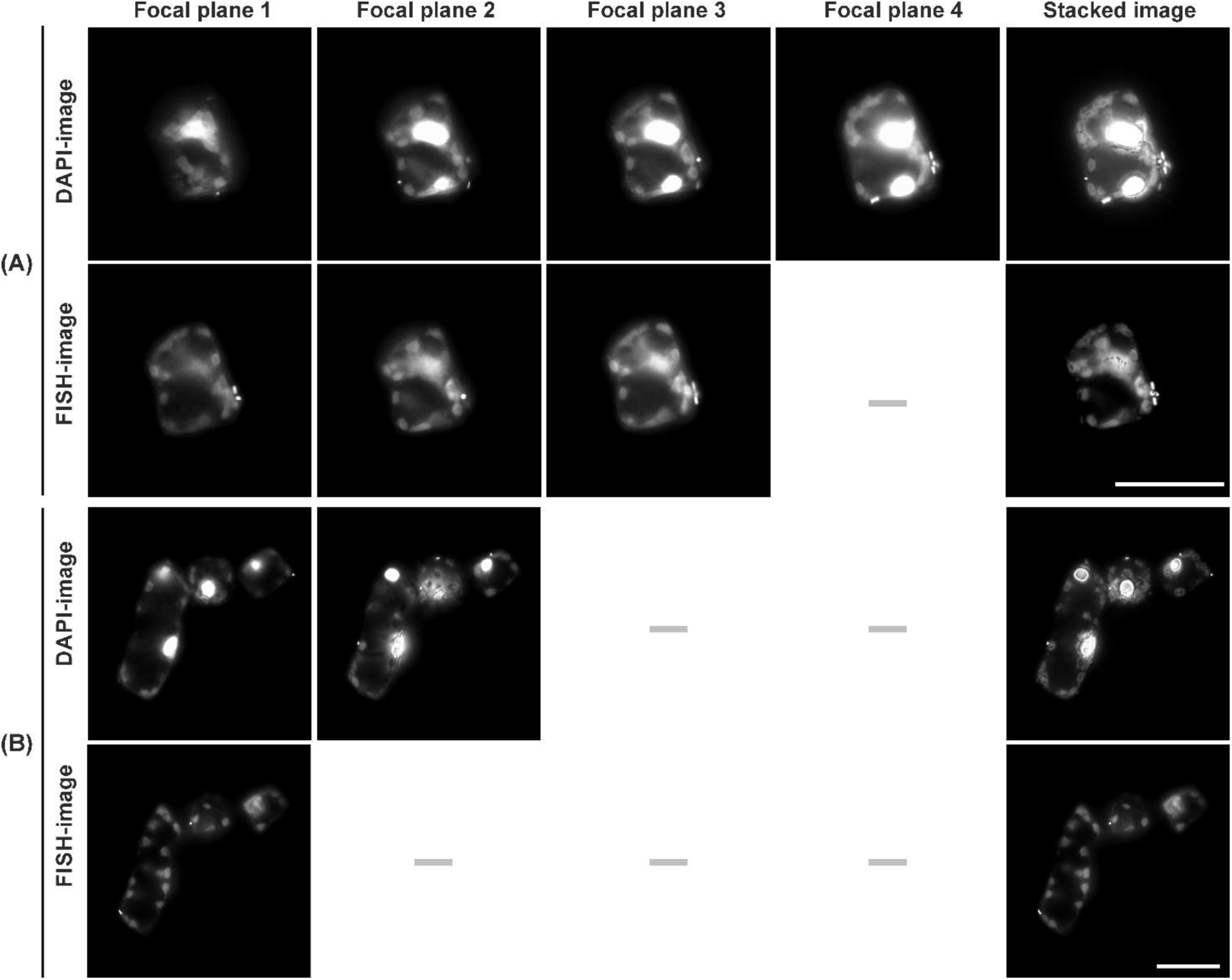
Raw images from different focal planes of Figures 3. The sign “–“ means that the image was not taken at the corresponding focal plane. Scale bar = 20 µm.

